# Development of a new flippase-dependent mouse model for red fluorescence-based isolation of Kras^G12D^ oncogene-expressing tumor cells

**DOI:** 10.1101/2024.07.26.605291

**Authors:** Dusan Hrckulak, Michaela Krausova, Jakub Onhajzer, Monika Stastna, Vitezslav Kriz, Lucie Janeckova, Vladimir Korinek

## Abstract

Proto-oncogene KRAS, GTPase (KRAS) is one of the most intensively studied oncogenes in cancer research. Although several mouse models allow for regulated expression of mutant Kras, selective isolation and analysis of transforming or tumor cells that produce the Kras oncogene remains a challenge. In our study, we present a knock-in model of oncogenic variant Kras^G12D^ that enables the “activation” of Kras^G12D^ expression together with production of red fluorescent protein tdTomato. Both proteins are expressed from the endogenous *Kras* locus after recombination of a transcriptional stop box in the genomic DNA by the enzyme flippase (Flp). We have demonstrated the functionality of the allele termed *RedRas* (abbreviated *Kras^RR^*) under *in vitro* conditions with mouse embryonic fibroblasts and organoids and *in vivo* in the lung and colon epithelium. After recombination with adenoviral vectors carrying the *Flp* gene, the *Kras^RR^* allele itself triggers formation of lung adenomas. In the colon epithelium, it causes the progression of adenomas that are triggered by the loss of tumor suppressor adenomatous polyposis coli (Apc). Importantly, cells in which recombination has successfully occurred can be visualized and isolated using the fluorescence emitted by tdTomato. Furthermore, we show that Kras^G12D^ production enables intestinal organoid growth independent of epidermal growth factor (EGF) signaling and that the Kras^G12D^ function is effectively suppressed by specific inhibitor MRTX1133.

## Introduction

Cancer is a complex disease characterized by uncontrolled proliferation and spread of abnormal cells. The transformation of normal cells into cancer cells is driven by sequential accumulation of genetic mutations that disrupt the normal regulatory mechanisms of cell growth, differentiation and apoptosis (Hanahan and Weinberg 2011).

One of the most common genetic alterations in cancer is the mutation of the *KRAS* gene (Kirsten rat sarcoma viral oncogene homolog) (Haigis 2017), a member of the Ras gene family. This family includes HRAS and NRAS; all RAS proteins have a highly conserved domain structure and are essential for the transmission of extracellular growth factor-derived signals to intracellular signaling networks. RAS proteins are small GTPases that switch between an inactive, GDP-bound state and an active, GTP-bound state. This transition is regulated by GDP–GTP exchange factors, which facilitate the exchange of GDP for GTP, and by GTPase-activating proteins (GAPs), which enhance the hydrolysis of GTP to GDP and return Ras to its inactive state. Mutated Ras proteins have impaired GTPase activity, which makes them resistant to inactivation by GAPs (Pylayeva-Gupta et al. 2011).

A significant proportion of human cancers have mutations in one of the *RAS* genes, with *KRAS* mutations being particularly common in certain cancers. In lung cancer, mutated *KRAS* is a hallmark of non-small cell lung cancer (NSCLC), particularly adenocarcinoma, where it promotes tumor development and progression (Ferrer et al. 2018). Similarly, *KRAS* mutations in colorectal cancer (CRC) contribute to tumor development (Cancer Genome Atlas 2012) and are associated with poor prognosis and resistance to epidermal growth factor receptor (EGFR)-targeted therapies (Lievre et al. 2006). In pancreatic ductal adenocarcinoma (PDAC), oncogenic *KRAS* is almost ubiquitously present and drives the development of these highly lethal tumors (Ryan et al. 2014).

Approximately 90 % of activating *KRAS* mutations occur in codons 12 and 13 of exon 1 of the *KRAS* gene and lead to an exchange of glycine for aspartate (G12D or G13D) or valine (G12V). Mutations in codon 61 of exon 2 account for about 5% of KRAS mutations and usually lead to the exchange of glutamine for arginine (Q61R), leucine (Q61L) and lysine (Q61K) (Hobbs et al. 2016; Yang et al. 2023). These mutations lead to continuous activation of the RAS/mitogen-activated protein kinase (MAPK) signaling pathway, which promotes cell proliferation (Bahar et al. 2023). The presence of *KRAS* mutations is often associated with aggressive tumor phenotypes and resistance to therapy (Bahrami et al. 2018; Ng et al. 2013), which underlines the urgent need for models that can accurately replicate these mutations for research and therapeutic development.

Site-specific recombinases have become indispensable tools for precise genetic manipulations in transgenic mouse models. The bacteriophage P1-derived Cre-loxP system uses Cre recombinase to catalyze site-specific recombination between loxP sites, which are 34-base pair sequences inserted at specific locations in the genome (Sauer and Henderson 1988). When two loxP sites are aligned in the same direction, Cre recombinase cuts out the DNA sequence between them, resulting in a deletion (Meinke et al. 2016).

The spatial specificity of Cre-mediated DNA recombination is achieved by using promoters/genes that control Cre expression in desired tissues or cell types. To enable temporal control over DNA recombination, the CreERT variant was developed. CreERT is a fusion protein consisting of Cre recombinase and a mutant human estrogen receptor ligand-binding domain (ERT) that prevents translocation of Cre to the nucleus in the absence of ligand (Feil et al. 1996).

An analogous approach for site-specific DNA rearrangements is derived from the yeast *Saccharomyces cerevisiae*. In this method, the enzyme flippase (Flp) is used to mediate recombination between two 34-base pairs of Flp recombinase recognition sites (FRTs) (O’Gorman et al. 1991). This system provides an additional level of genetic control and allows sequential gene activation or deletion, especially when used in conjunction with the Cre-loxP system. To address the thermal instability of Flp and its low recombination efficiency in mammalian cells, an enhanced version of the enzyme called Flpe was produced that has higher activity at physiological temperatures (Buchholz et al. 1998). Subsequently, a codon-optimized version called Flpo, was developed (Raymond and Soriano 2007). This variant improves both the efficiency and stability of recombination, making it more suitable for complex genetic engineering applications in mammalian cells.

Both the Cre-lox and Flp-FRT systems have been used to study the proto-oncogene *Kras* in mice (Jackson et al. 2001; Tuveson et al. 2004; Young et al. 2011). In our study, we present the generation of a new mouse strain that activates the oncogenic variant Kras^G12D^ in the target tissue after induction by the Flp-FRT system. Importantly, the cells producing Kras^G12D^ simultaneously begin to express a tandem dimer variant of red fluorescent protein Tomato (tdTomato), which is known for its brightness and photostability (Shaner et al. 2005). This protein enables the isolation and subsequent analysis of these (transforming) cells. The study includes a series of tests that demonstrate the functionality of the new *RedRas* (*Kras^RR^*) allele.

The described mouse model will be a suitable tool for analyzing the efficacy and specificity of potential drugs targeting different oncogenic variants of the Kras protein. Among other things, the design of the allele allows it to be combined with a number of mouse models based on Cre-loxP recombination to model the stepwise accumulation of mutations during tumor progression.

## Materials and methods

### Gene targeting in embryonic stem (ES) cells

A Flippase-removable stop box was generated as previously described (Kasparek et al. 2014). Briefly, a neomycin resistance expression cassette consisting of the mouse phosphoglycerate kinase 1 (*pGK*) promoter followed by aminoglycoside 3‘-phosphotransferase and two consecutive herpes simplex virus thymidine kinase (*HSV TK*) and simian virus early (*SV40E*) polyA signals was used as a transcriptional stop box. This element was flanked by a pair of consensus FRT sites, followed by a chimeric chicken actin β and a rabbit β-globin splice acceptor from the pCAGEN vector scaffold (kindly provided by Connie Cepko via Addgene; #13775). The cDNA of the mouse *Kras^G12D^*variant (kindly provided by Sandra Orsulic via Addgene; #11549) and the cDNA of *tdTomato* (a gift from Roger Tsian, University of California, San Diego, CA, USA) were each cloned into the pIRES vector (Takara Bio, Japan) to assemble a bicistronic coding sequence. Both homology arms were derived from the mouse 129 genomic fragment in the BAC clone bMQ415f20. The short homologous arm was derived from intron 1 of the *Kras* gene using the forward primer 5’-ATTGCCGCGGCCACAAAGGAGGAGGT-3’ and the reverse primer 5’-CGCGCGCGTTTAAACGTAAAACTCTAAGATATTCC-3’ for amplification and the SacII and PmeI restriction endonuclease sites for cloning. The long homologous arm was derived from intron 2 of the *Kras* gene using the forward primer 5’-AAAGGTCGACAGCCAGCCGCTTTGA-3’ and the reverse primer 5’-AATATCCCGGGAATCATTGCTGCAATC-3’ for amplification and the Sall and SmaI restriction endonuclease sites for cloning.

The targeting construct was incorporated into the pGKneoF2L2DTA vector (kindly provided by Philippe Soriano via Addgene; #13445). ES R1 cells were grown on a layer of mouse embryonic fibroblast (MEF) feeder cells (Stem Cell Technologies, CA, US) treated with mitomycin C (for 2 hours at a final concentration of 10 μg/mL; purchased from Merck, NJ, US). ES cells were cultured in Glutamax Dulbecco’s modified Eagle’s medium (DMEM; Thermo Fisher Scientific, MA, US) supplemented with 15% fetal bovine serum (FBS; ES cells tested; Thermo Fisher Scientific), 2 mM l-glutamine, 1 mM sodium pyruvate, 1× non-essential amino acids, 0.1 mM β-mercaptoethanol, 100 UI penicillin/streptomycin (all chemicals were purchased from Thermo Fisher Scientific). The complete medium was supplemented with conditioned media obtained from COS-7 cells (kindly provided by Vladimir Divoky; Palacky University, Olomouc, Czech Republic) stably expressing murine leukemia inhibitory factor (LIF). The targeting vector (24 μg) linearized with SacII restriction endonuclease was electroporated into 1 × 107 ES R1 cells using the Gene Pulser II system (Bio-Rad, CA, US) (settings: 380 V, 25 μF, time constant ∼3.4 s). Cells carrying the integrated construct were selected with G418 (350 µg/mL, Gibco). After 7 days, genomic DNA (gDNA) isolated from 119 selected clones was analyzed by PCR for the presence of short and long homologous arms. gDNA from seven positive clones was digested with restriction enzyme SexAI and analyzed by Southern blotting for successful homologous recombination on both arms of the target construct. A template for probes was produced by PCR using primers 5’-GGCAAGCTTCCTGGTGCTGGGAGG-3’ and 5’-GCGGCTCGAGCACAGCCCTTACAGG-3’ for a short-arm probe and primers 5’-CACAAAGCTTACGTTGGAAATGTTG-3’ and 5’-CTGAGCTCGAGAAAAGAACAAAATA-3’ for a long-arm probe. Correctly targeted ES cell clones were karyotyped, and cells of clones #10 and #26 with the correct chromosome number were injected into blastocysts from superovulated C57BL/6J females. Blastocyst injections were performed with the help of the Transgenic Unit (Institute of Molecular Genetics, Prague, Czech Republic). Out of five chimeric males, one male was mated with C57BL/6J females to generate an F1 generation of *Kras^wt/Flip-ready^*heterozygous mice.

### Mouse strains and genotyping

The production, housing of mice and *in vivo* experiments were performed in accordance with the Council Directive of the European Communities of November 24, 1986 (86/609/EEC) and national and institutional guidelines. Animal care and experimental procedures were approved by the Animal Care Committee of the Institute of Molecular Genetics (approval numbers 63/2019, AVCR 6566/2022 SOV II). Genotyping of animals was performed using lysates from tail tip biopsies as previously described (Pospichalova et al. 2011). For genotyping of *Kras* wild-type (wt) and targeted *Kras^Flp-ready^* alleles, the following primers were used: common forward primer P1: 5’-GTCGACAAGCTCATGCGGG-3’, wt reverse primer P2: 5’-CATGTCTCACACAAGATTATCAAAA-3’ and *Kras^Flp-ready^* reverse primer P3: 5’-CGAAGGAGCAAAGCTGCTA-3’. PCR with the wt and *Kras^Flp-ready^*alleles yielded 325-bp and 532-bp DNA fragments, respectively. Examples of genotyping results are shown in Fig. 1. Successful recombination of the *Kras^Flp-ready^* allele was verified with the forward primer P4: 5’-GGGTGTGTCCACAGGGTATAGCGTACTATG-3’ and the reverse primer P5: 5’-TCAGTCATTTTCAGCAGACCGGTCCG-3’, resulting in a 447-bp DNA fragment in *Kras^RR^*cells. The mouse strains *ACTB-FLPe* (Rodriguez et al. 2000)(full name of strain B6.Cg-Tg(ACTFLPe)9205Dym/J) and *Rosa26-STOP-tdTomato* mice (Madisen et al. 2010) (strain 6;129S6-Gt(ROSA)26^Sortm14(CAG-tdTomato)Hze/J^) were purchased from The Jackson Laboratory (Bar Harbor, ME, US). *Apc^cKO/cKO^* mice (Kuraguchi et al. 2006) (strain B6.Cg-Apc^tm2Rak^) were purchased from the Mouse Repository (National Cancer Institute, Frederick, MD, US). Animals were maintained under specific pathogen-free conditions and genotyped using tail biopsies according to protocols provided by the suppliers.

**Figure 1.**
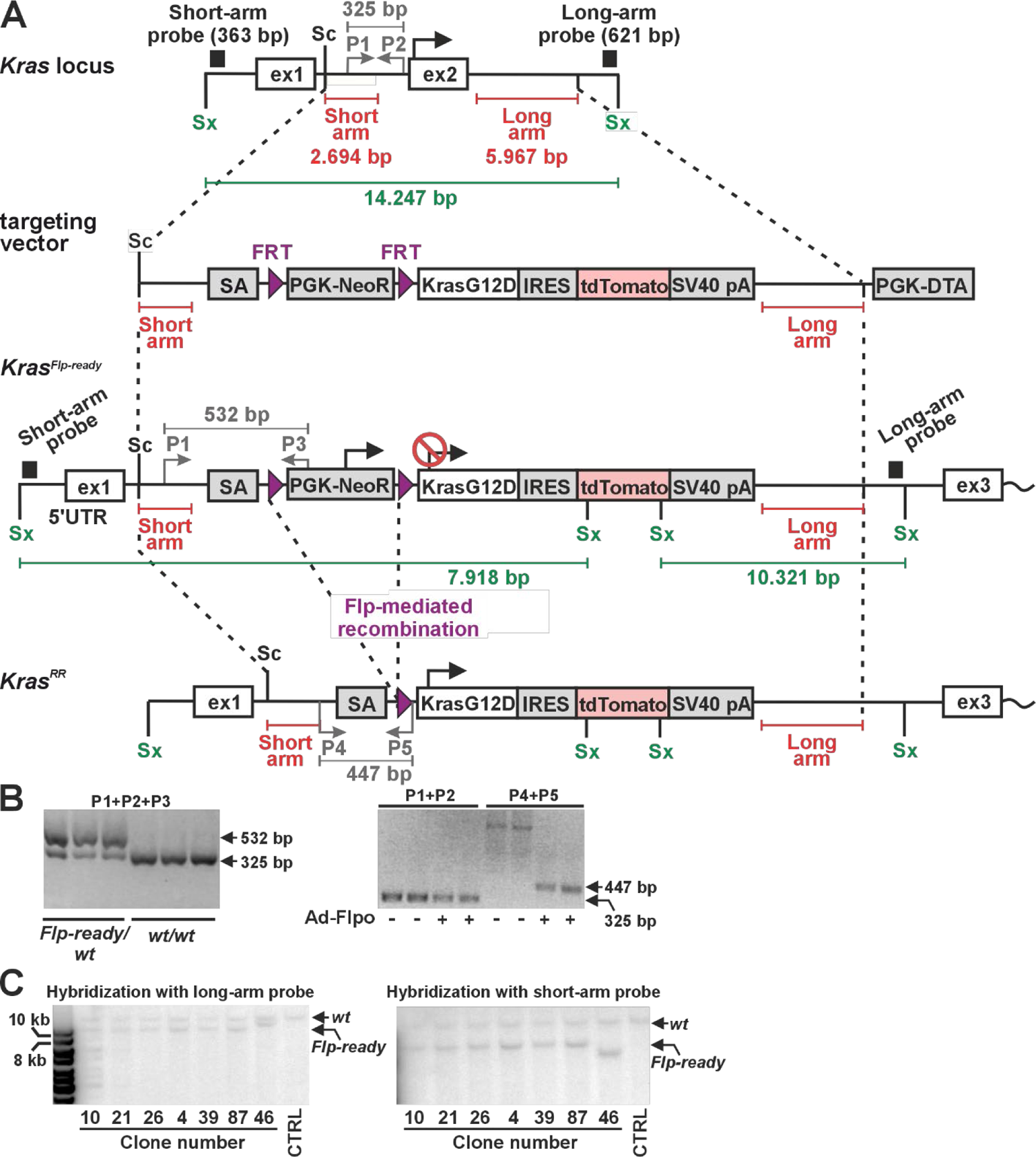
Generation of mice expressing an oncogenic Kras^G12D^ variant and the red fluorescent protein tdTomato after flippase (Flp)-mediated recombination of the transcription blocker. **a** Schematic representation of *Kras* locus targeting by homologous recombination in mouse embryonic stem (ES) cells. The diagram shows the *Kras* locus before and after targeting and the main components of the targeting vector. The parts of the locus that are present in the targeting vector, i.e., the short and long arms, are indicated by red lines. The G418 resistance expression cassette (PGK-NeoR) serves as a transcriptional blocker that, in combination with the β-globin splice acceptor (SA) inserted into the first intron of the *Kras* gene, terminates transcription from the endogenous *Kras* promoter and effectively generates a *Kras* null allele, *Kras^Flp-ready^*, which can be activated after Flp-mediated excision of the blocker. The lower diagram shows the situation after Flp-mediated recombination of DNA flanked by the Flp recombinase recognition sites (FRT; purple triangles). The allele producing *Kras^G12D^*-internal ribosomal entry site (IRES)-tdTomato mRNA was designated *Kras^RR^*. Two exons of *Kras*, where the first exon contains the 5’ untranslated region and the Kras protein is translated from the second exon, are indicated by black recangles. The translation start site (black arrow) is active in the *Kras^RR^* allele. The position of the probes used to verify the correct targeting of the locus at the 5’ (short arm probe) and 3’ (long arm probe) ends of the long construct is indicated by black rectangles. P1, P2 and P3 indicate the position of primers used for genotyping of the *Kras^wt^* (P1 and P2) or *Kras^Flp-ready^* allele (P1 and P3). The size of the bands obtained by Southern blot analysis after probing genomic DNA digested with restriction endonuclease SexAI (Sx) is indicated by green lines. Sc: SacII restriction endonuclease site. PGK-NeoR: the sequence of the mouse phosphoglycerate kinase 1 (PGK) promoter that drives the expression of the neomycin kanamycin phosphotransferase type II gene (NeoR), terminated by the thymidine kinase of herpes simplex virus polyadenylation signal (pA) followed by pA of simian virus 40 (SV40 pA; pAs are not indicated in the scheme); PGK-DTA: an expression cassette for counter-selection of ES cell clones with random insertion of the target construct. The cassette contains PGK, which drives expression of the diphtheria toxin A (DTA) chain terminated by SV40 pA (not shown). For clarity, the size of the symbols for each genetic element is not proportional to the length of its actual sequence. **b** Left, an example of PCR-based genotyping of ES cell clones of the indicated genotype. A mixture of P1, P2 and P3 primers was used. Right, PCR detection of the *Kras^RR^* allele in genomic DNA isolated from intestinal organoids derived from *Kras^wt/Flp-ready^*mice. Some organoids were infected with Flpo-expressing adenoviruses (Ad-Flpo) as indicated and analyzed for the presence of the *Kras^wt^*(primers P1+P2) or *Kras^RR^* (primers P4+P5) allele. **c** Southern blot analysis to verify the correct genomic integration of the targeting vector. The blot shows the hybridization results with the long- and short-arm probes confirming the presence of the expected recombination events at the 5’ and 3’ ends of the targeting locus in selected ES cell clones. Genomic DNA was cut with restriction endonuclease SexAI; CTRL, line with genomic DNA from non-targeted ES cells. The DNA marker is shown on the left.

### Isolation of MEFs and *in vitro* recombination

Mouse embryonic fibroblasts were isolated from day 12.5 (E12.5) embryos as previously described (Janeckova et al. 2015). The dissected embryos were washed in cold phosphate-buffered saline (PBS), and the head and internal organs (heart, liver) were removed and used for genotyping. The remaining embryonic tissues were minced, incubated in trypsin/EDTA solution (15 minutes at 37°C; Sigma), further dissociated by vigorous pipetting, and then cultured in Iscove’s Modified Dulbecco’s Medium (IMDM; Gibco) supplemented with 10% fetal bovine serum (FBS), penicillin/streptomycin (100 UI), 2 mM L-glutamine and 1× non-essential amino acids (all reagents from Gibco). *Kras^RR^* expression was induced *in vitro* by adenovirus-mediated recombination. *Kras^wt/Flip-ready^*MEFs at 70% confluence were inoculated with Ad-FLPo (Vector Biolabs, PA, US) at multiplicity of infection (MOI) of 1000 and incubated overnight. Similarly, tdTomato expression was induced in *Rosa26-STOP-tdTomato* MEFs by infection with Ad-Cre (Vector Biolabs). After infection, cells were analyzed by live cell imaging using a Leica Stellaris confocal microscopy system (Leica, Germany) equipped with a 63 × water immersion objective. The resulting images were deconvoluted using Huygens software (SVI, Netherlands). Alternatively, cells were analyzed by flow cytometry as described in the following sections.

### Flow cytometry

#### Analysis of MEFs and lung epithelium

Mouse embryonic fibroblasts were harvested with trypsin-EDTA solution (Merck), washed with complete growth medium and pelleted. The cell pellet was resuspended in PBS, and the cell suspension was filtered through a 40-µm cell strainer (Corning, NY, US). Cells were then treated with Hoechst 33258 (Thermo Fisher Scientific) to distinguish live from dead cells. Live cells were analyzed for endogenous fluorescence using an LSRII flow cytometer (BD Biosciences, NJ, USA). For analysis of the lung epithelium, mice were euthanized by cervical dislocation. The thorax was opened and the lungs were perfused with 5 ml PBS with Dispase I (Thermo Fisher Scientific; stock solution 100 mg/ml, diluted 1:300) through the right ventricle. The lungs were then excised, rinsed externally with PBS on a Petri dish, incubated at room temperature for 20 minutes, and cut into pieces. These pieces were digested with Dispase I and DNAse I (Thermo Fisher Scientific; working concentration 1 U/ml) in serum-free IMDM medium for three 10-minute cycles at 37°C and 800 rpm on a rotating platform. After each incubation, the resulting cell suspension was harvested into cold IMDM supplemented with 10% FBS (Gibco), and fresh cleavage medium was added to the remaining tissue. The cells were then filtered through a 40 µm cell strainer and pelleted. The cell pellet was thoroughly resuspended, treated with ACK lysis buffer (Gibco) for 5 minutes on ice to reduce red blood cell contamination, and then washed with PBS. Cells were stained with Pacific Blue™ (PB)-conjugated anti-CD45 antibody (#103126, BioLegend, CA, US; dilution 1:200), PB-conjugated anti-CD31 antibody (#102422, BioLegend; 1:200), and fluorescein (FITC)-conjugated anti-EpCAM antibody (#11-5791-82, Thermo Fisher Scientific; 1:400) for 20 minutes at 4°C. Cells were sorted by forward scatter (FSC), side scatter (SSC), and negative staining for Hoechst and PB. Individual EpCAM^+^ cells were analyzed for endogenous tdTomato fluorescence using an Influx cell sorter (BD Biosciences).

#### Intestinal organoids

Intestinal organoids were harvested by pipetting, washed with PBS and processed into a single cell suspension by incubation with TrypLE Express (Thermo Fisher Scientific) at 37 °C for 15 minutes. The cells were then washed with IMDM supplemented with 10 % FBS, filtered through a 40 µm cell strainer and pelleted. The cell pellet was resuspended in fresh PBS and treated with Hoechst 33258 to identify viable cells. The endogenous fluorescence of tdTomato was analyzed using an LSRII flow cytometer.

### Adenovirus-mediated *in vivo* recombination

Replication-deficient adenoviruses expressing Cre, FLPo and CreERT2-P2A-FLPo (Ad-Cre, Ad-FLPo, Ad-CreERT2-P2A-FLPo) were obtained from Vector Biolabs. For *in vivo* applications, adenovirus stocks were propagated by repeated infections of HEK293 cells (Cat. No.: CRL-1573, American Type Culture Collection, VA, US) and concentrated using the AdEasy Virus Purification Kit (Agilent, CA, US) according to the manufacturer’s instructions to achieve a concentration of up to 3 × 10^12^ plaque-forming units (PFU)/ml in storage buffer under BSLII conditions. Quantification of intact viral particles was performed by flow cytometry on MEFs derived from Rosa26-STOP-tdTomato (adenoviruses producing Cre; Ad-Cre) and Rosa26-STOP-Blueflirt (Kasparek et al. 2014) (adenoviruses producing Flpo; Ad-Flpo) compared to the manufacturer’s original stocks and by plaque assays in HEK293 cells. Notably, we found that CreERT2 recombinase in the latter adenovirus acted independently of 4-hydroxytamoxifen during the titrations.

To initiate Kras^RR^-dependent transformation of the airway epithelium, intranasal infection was performed as previously described (DuPage et al. 2009). In brief, 5 × 10^7^ PFU of Ad-FLPo in 75 µl PBS were administered dropwise into a single nostril of 10-week-old KrasFlip-primed mice anesthetized with isoflurane (Baxter, IL, US). Animals were sacrificed 8 or 12 weeks post-infection for histologic examination or flow cytometric analysis of epithelial cells. Ad-Cre or Ad-CreERT2-P2A-FLPo was administered rectally to transform the colonic epithelium. Ten-week-old *APC^cKO/cKO^/Kras^Flip-ready^* animals were anesthetized with isoflurane and their colons were irrigated with 50% ethanol in 100 µl PBS using a soft cannula to disrupt the mucus barrier. After a 5-hour recovery period, 1 × 10^9^ PFU of adenovirus was injected rectally with a soft cannula into the isoflurane-anesthetized animals, which were placed on a tiltable surface 5 minutes after injection to prevent leakage. Animals injected rectally with the virus were killed 10-12 weeks later and their colons were histologically examined for adenoma formation.

### Histology, immunohistochemical staining and fluorescence microscopy

Mouse lungs and small intestine were fixed overnight in 10 % neutral buffered formaldehyde in PBS and embedded in paraffin using an automated tissue processor (Leica, Germany). Sections of 5 µm were stained according to the protocol published by Hrckulak et al. (Hrckulak et al. 2018). In brief, samples were deparaffinized in xylene and stained with hematoxylin and eosin (Vector Laboratories) according to the manufacturer’s instructions.

For immunohistochemical staining, antigen retrieval was performed in a steam bath by immersing the slides in 10 mM citrate buffer, pH 6.0. Endogenous peroxidase activity was blocked by incubating the slides in 0.2 % H_2_O_2_ (Merck; strain 30 %) in methanol (Merck) for 20 min. Samples were incubated overnight at 4 °C with the primary anti-PCNA antibody (ab18197, Abcam, Cambridge, UK). A biotin-conjugated secondary anti-rabbit antibody was then applied (Biotin-XX Goat anti-Rabbit IgG, B-2770, Thermo Fisher Scientific). The peroxidase signal was enhanced by a 30-minute incubation with the Vectastain ABC kit (Vector Laboratories) and developed with DAB solution (Vector Laboratories). Samples were counterstained with hematoxylin, and brightfield images were taken using a DM6000 microscope (Leica).

For fluorescence microscopy, the primary antibodies anti-E-cadherin (ab231303, Abcam) and anti-Trop2 (ab214488, Abcam) were used together with secondary Alexa488-conjugated goat anti-mouse and Alexa594-conjugated goat anti-rabbit antibodies (Thermo Fisher Scientific). Cells were counterstained with DAPI nuclear stain (Sigma-Aldrich). Microscopic images were captured with the Stellaris confocal system (Leica). Images were processed using Huygens software (SVI, NE) and the Fiji package (Schindelin et al. 2012).

### Organoid cultures, *in vitro* recombination, live cell imaging, and proliferation assay

Organoid cultures were prepared from freshly isolated intestinal crypts according to the methods previously described by (Sato et al. 2011; Sato et al. 2009). For this purpose, the middle section of the small intestine was cut longitudinally and the villi were scraped off with a coverslip. The tissue was then washed several times in PBS. The tissues were incubated with a 5 mM EDTA pH 8 solution (Merck) at 4°C for 30 minutes, followed by gentle shaking to detach the remaining differentiated epithelial cells. The tissue was then transferred to fresh PBS and shaken vigorously to release the crypts into the supernatant. The supernatant containing the crypts was filtered through a 70μm strainer (Corning).

The pelleted crypts were centrifuged at 180 × g for 5 min at 4 °C, washed in advanced DMEM/F-12 medium (Thermo Fisher Scientific), and embedded in Matrigel (BD Biosciences). Embedded crypts were maintained in advanced DMEM/F-12 medium supplemented with 1 × N-2 supplement, 1 × B-27 supplement (both from Thermo Fisher Scientific), mRspo1 (500 ng/mL; Peprotech, Rocky Hill, NJ, USA), mNoggin (100 ng/mL; Peprotech), 10 mM 4-(2-hydroxyethyl)-1-piperazineethanesulfonic acid (HEPES; Thermo Fisher Scientific), 1 × Glutamax, 1 mM N-acetylcysteine (Sigma-Aldrich), 1 × penicillin/streptomycin, mouse epidermal growth factor (mEGF, 50 ng/mL; Thermo Fisher Scientific), and primocin (100 µg/mL, Invivogen, Toulouse, France). This media composition is referred to as ENR.

Kras^RR^ expression in *Kras^Flip-ready^* organoids was induced by *in vitro* adenoviral recombination. Organoids cultured for 72 hours in ENR media supplemented with recombinant Wnt surrogate FC fusion protein (5 nM, Thermo Fisher Scientific), Rho-associated protein kinase (ROCK) inhibitor Y-27632 (2.5 mM, Merck) and nicotinamide (10 mM, Merck) were harvested by pipetting Matrigel. They were then washed with PBS and processed into cell clusters by incubation with TrypLE Express (Thermo Fisher Scientific) at 37 °C for 5 min. The cell clusters were spinnoculated with Ad-FLPo at 1000 MOI at 600 × g for 60 minutes at 30°C and then incubated for 6 hours at 37 °C, 5% CO2 in WENRY+NIC media. The cell clusters were then washed with PBS, resuspended in Matrigel and cultured in the same media until the organoids were reconstituted. After reconstitution, the organoids could be further cultivated in ENR media.

NR medium (without mEGF) with or without EGFR tyrosine kinase inhibitor gefitinib (5 μM, MedChemExpress, NJ, US) was used to select cells independent of EGF-EGFR signaling. MRTX1133 (100 nM, MedChemExpress) was used to specifically inhibit Kras^G12D^ (Kras^RR^) activity in cultured cells. The selected organoids were imaged with brightfield microscope EVOS XL Core (Thermo Fisher Scientific) over 10 days of cultivation and one passage.

To evaluate the growth of organoids under selective conditions, we also used the xCELLigence RTCA eSight system (Agilent) with a 10 × objective. The Immune Cell Clustering and Proliferation module was used for automatic focus adjustment and stitching of two images per well for brightfield imaging. Organoids were seeded in a 96-well plate (Corning) in eight technical replicates, and each well was imaged every 4 hours in the appropriate growth media (control samples infected with Ad-Cre in ENR, other samples infected with Ad-Flpo in NR+ gefitinib). After 72 hours, the media were changed, selection was performed, and imaging continued for the next 90 hours. The original RTCA software was used to automatically segment the organoid area in each field of view and to calculate the average area for all samples. Automatic segmentation was optimized using training images for all acquired time points. Data were normalized to the 72-hour time point, and the time dependence of relative organoid area was recorded.

### RNA isolation, reverse transcription quantitative PCR (RT-qPCR)

Total RNA was isolated from extraembryonic tissues of partially resorbed Kras^wt/*RR*^ embryos using TRI reagent (Merck), while RNA was extracted from intestinal organoids using the RNeasy Mini Kit (Qiagen, DE). For cDNA synthesis, 1 µg of total RNA was used with random hexamer primers in a reaction volume of 20 µl. The RNA was reverse transcribed with RevertAid Reverse Transcriptase (Thermo Fisher Scientific) according to the manufacturer’s protocol. *Kras* cDNA was amplified for sequencing using non-selective primers covering exons 1, 2 and 3 of both Kras^wt^ and Kras^RR^ transcripts (forward primer 5’-ATTTCGGGACCCGGAGCGA-3’ and reverse primer 5’-CCCTCCCCAGTTCTCATGTA-3’). This was performed with the EliZyme HS Robust Mix (Elisabeth Pharmacon, CZ). The PCR products were then subjected to Sanger sequencing with SEQme (CZ) from the reverse primer. Quantitative RT-PCR was performed in triplicate using the SYBR Green I Master Mix in a LightCycler 480 system (Roche, CH) and the following primers: mPhdla1 forward 5’-CTCACCCGTACCAACTCCAG-3’, reverse 5’-GTGGGGAGACTCTGTTGGTT-3’; mSerpinb1a forward 5’-ATGCTCCATTCCGACTGAGT-3’, reverse 5’-CTTAAGACCCGTGGACTCGT-3’; mEtv4 forward 5’ ACCTTAGTTGTGTCATCCCCC--3’, reverse 5’-CATTTCCGGGAGATTTGCTGC-3’; mEtv5 forward 5’-AGGAGCCCCGAGATTACTGT-3’, reverse 5’ CTCGGGTACCACGCAAGTAT--3’; mDusp4 forward 5’-GTGAGCCTGGAGCAGATCCT-3’, reverse 5’-CCACCTTTAAGCAGGCAGAT-3’; mDusp6 forward 5’-TTGAATGTCACCCCCAATTT-3’, reverse 5’-GCTGATACCTGCCAAGCAAT-3’; mKi67 forward 5’-GTAACCTGCCTGCGAAGAGA-3’, reverse 5’-TTTCGCAACTTTCGTTTGTG-3’; mUbb forward 5’-ATGTGAAGGCCAAGATCCAG-3’, reverse 5’-TAATAGCCACCCCTCAGACG-3’; and mActb forward 5’-GATCTGGCACCACACCTTCT-3’, reverse 5’-GGGGTGTTGAAGGTCTCAAA-3’.

## Results and Discussion

### Production of mice expressing the Kras^G12D^ oncogene and tdTomato from the endogenous *Kras* locus

For historical reasons, most conditional alleles of investigated genes, including tumor suppressors and oncogenes, are based on the Cre-loxP system. To investigate the influence of oncogenic activation of the *Kras* gene on the development and progression of neoplasms, we created a new *Kras* allele that activates the oncogenic variant *Kras^G12D^* after removal of two FRT sites by Flp-mediated recombination. For subsequent analysis of cells expressing *Kras^G12D^*, we also inserted a sequence encoding the red fluorescent protein tdTomato into the *Kras* locus. This protein is translated together with Kras^G12D^ from a single bicistronic mRNA containing an internal ribosomal entry site (IRES) between the *Kras^G12D^* and *tdTomato* sequences. This mRNA is transcribed from the *Kras* locus after removal of a transcriptional blocker flanked by FRT sites. This blocker, which consists of an expression cassette that produces neomycin phosphotransferase and induces resistance to G418 in ES cells, was inserted into the first intron of the *Kras* gene. The targeting strategy is shown in Fig. 1A. The targeting construct was electroporated into ES cells; 119 G418-resistant ES cell clones were expanded and analyzed by PCR. Seven of these clones were further analyzed by Southern blotting, six of which confirmed correct integration at both the 5’ and 3’ ends of the construct Fig. 1B. The resulting clones were karyotyped, and two clones (nos. 10 and 26) were injected into blastocysts. From a total of seven chimeric mice, three founder animals were obtained that were heterozygous for the conditional *Kras* allele; this allele was designated *Kras^Flp-ready^* (Fig. 1C). These animals were used for subsequent analyzes.

Genotyping of the progeny revealed that no viable mice expressed the *Kras^G12D^-IRES-tdTomato* mRNA, which begins to be produced after Flpe-mediated recombination of the *Kras^Flp-ready^* allele. The resulting allele was named *Kras^RR^*. Analysis of the embryos at embryonic day 9.5 (E9.5) showed that the *Kras^wt/RR^* embryos were smaller and paler than the *Kras^wt/wt^* embryos. The mutant embryos were clearly unable to develop further; analysis at later stages showed partially resorbed embryos (Fig. 2A). Subsequent cDNA analysis revealed that the resorbed embryos expressed both the wt and oncogenic *Kras^G12D^* variants, while healthy embryos only produced mRNA encoding wt *Kras* (Fig. 2B). The phenotype described above essentially confirmed the results of Tuveson and colleagues obtained with a similarly constructed *Kras^LSL-G12D^* allele, but using the Cre-loxP system activated by a *CMV-Cre* driver that is active in all cells of the mouse embryo (Tuveson et al. 2004). We then produced mouse embryonic fibroblasts (MEFs) from *Kras^wt/Flp-ready^* embryos that emitted red fluorescence after infection with an adenovirus producing Flpo recombinase (Ad-Flpo). Control fibroblasts were obtained from mouse embryos carrying the ‘reporter’ allele *R26-stop-tdTomato* (hereafter *R26-tdTomato*), in which a transcriptional blocker flanked by loxP sites was inserted into the ubiquitously active *Rosa26* locus, followed by the *tdTomato* gene (Madisen et al. 2010). When these control fibroblasts were infected with an adenovirus that produces Cre recombinase (Ad-Cre), the blocker sequence was removed and the cells began to produce tdTomato, similar to cells of the *Kras^wt/RR^*genotype. We then performed FACS analysis on MEFs derived from *Kras^wt/Flp-ready^* mice. Some of these cells were control cells, while others were infected with Ad-Flpo. As shown in Fig. 2D, fluorescence-activated cell sorting (FACS) analysis revealed that a significant proportion of cells infected with Ad-Flpo emitted a red fluorescent signal.

**Figure 2.**
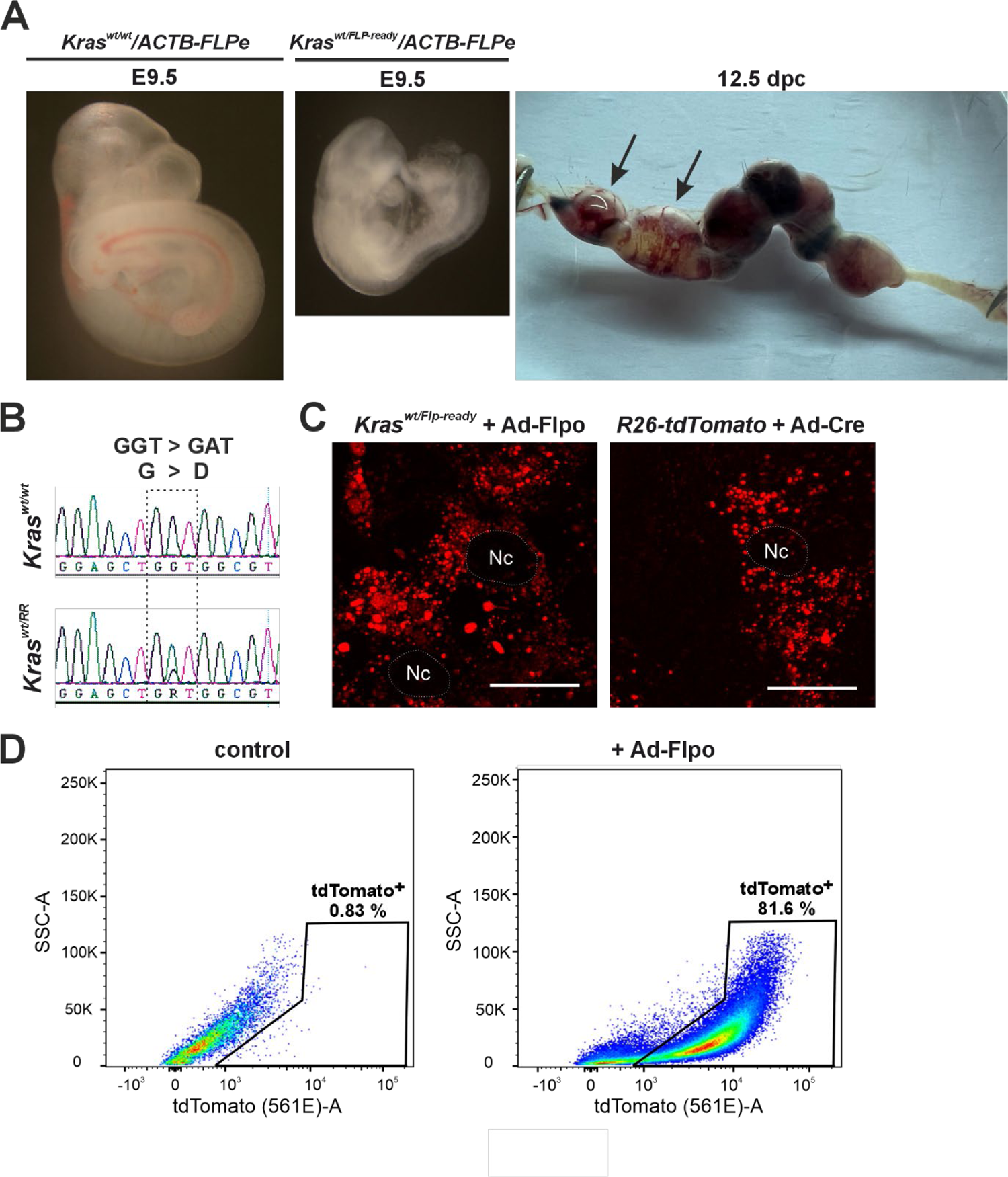
Analysis of the *Kras^RR^* allele during embryogenesis and in embryonic mouse fibroblasts (MEFs) **a** Left, brightfield micrographs of embryos with the indicated genotypes, isolated at embryonic day 9.5 (E9.5). Right, uterus isolated from an *ACTB-FLPe* female crossed with a *Kras^wt/FLP-ready^* male 12.5 days post-coitum (12 dpc). Black arrows indicate (partially) resorbed embryos. **b** Messenger RNA expressed from the oncogenic allele *Kras^RR^* is detected in the amniotic sacs of resorbed embryos. Sequencing of cDNA isolated from total RNA from the amniotic sac of healthy (upper diagram) or resorbed (lower diagram) embryos shows a single nucleotide substitution in codon 12 (GGT > GAT) of exon 1 (dashed frame). This substitution, in which glycine is converted to aspartate (G > D), only occurs in resorbed embryos and is an indication of the presence of the *Kras^RR^* allele. **c** Fluorescence micrographs of MEFs isolated from *Kras^wt/Flp-ready^* and *R26-tdTomato* animals. Images were taken 48 hours after infection with adenoviral particles expressing mouse codon-optimized Flp (Ad-Flpo) or adenoviral particles expressing Cre (Ad-Cre); in both cases, Flpo- or Cre-mediated recombination removes the “STOP box” and activates expression of the fluorescent reporter protein tdTomato (red fluorescent spots). The dashed areas show the cell nuclei (Nc). Scale bar: 10 µm. **d** Representative diagrams show fluorescence-activated cell sorting (FACS) of MEFs isolated from *Kras^wt/Flp-ready^*embryos after infection with “empty” adenoviral control particles (left) or Ad-Flpo (right). Viable (Hoechst 33258-negative) cells were visualized using the side scatter - pulse area (SSC-A) and endogenous tdTomato fluorescence; tdTomato-positive (tdTomato^+^) cells are shown in the black frame.

### The oncogenic *Kras^RR^* allele causes lung tumors

Recombinant adenoviruses are effective tools for ensuring expression of exogenous genes in target cells. Adenoviruses can infect a broad spectrum of cells and do not integrate into the host genome; therefore, the expression of the transferred gene is transient and does not lead to insertional mutations in the genome of the infected cell (Amalfitano 2004). As mentioned in the introduction, oncogenic mutations in *Kras* are found in at least a quarter of human lung adenocarcinomas. Instillation of adenoviruses has previously been used for Cre- and Flpe-dependent activation of oncogenic *Kras* alleles to induce lung tumors (Jackson et al. 2001; Tuveson et al. 2004; Young et al. 2011). To determine whether our prepared allele was functional, *Kras^wt/Flp-ready^*mice were infected with the adenovirus Ad-Flpo. Consistent with published results, we observed the formation of lung lesions approximately one month after instillation. Figure 3A shows a representative histologic image of a *Kras^wt/Flp-ready^* mouse 8 weeks after Ad-Flpo infection and the subsequent enlargement of these lesions 12 weeks after instillation. Of note, Ad-Flpo infection alone did not induce lesions in wt mice (Fig. 3A, left).

**Figure 3.**
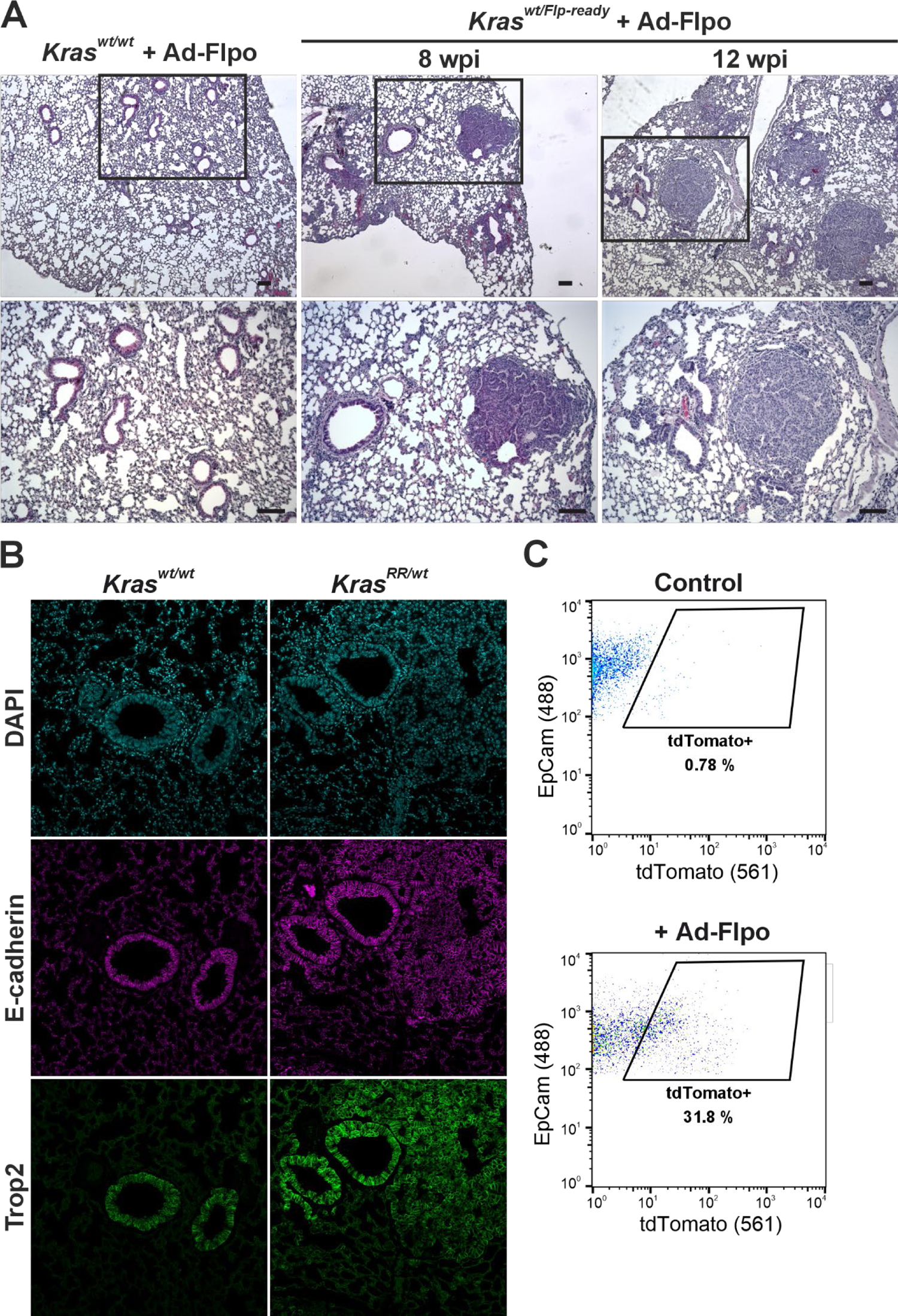
Tumor formation in the lung expressing the *Kras^RR^* allele **a** Histologic analysis of lungs isolated from *Kras^wt/Flp-ready^* animals 8 and 12 weeks post-infection (wpi) with Ad-FLPo particles (middle and right panels); lungs from infected *Kras^wt/wt^* mice (left panel) served as controls. In the lungs of *Kras^wt/Flp-ready^* mice, lesions adjacent to the airways were detectable as early as 8 weeks post-infection, which were absent in the controls. The images below show greater magnification of the framed areas. Sections were stained with hematoxylin nuclear stain (blue) and eosin cytoplasmic stain (pink). Scale bar: 50 µm. **b** Fluorescence micrographs showing the localization of E-cadherin (magenta) and trophoblast cell surface antigen 2 (Trop2, green) in the lungs of *Kras^wt/wt^* and Kras*^wt/RR^* mice 12 weeks after Ad-Flpo infection. The images show high expression of both proteins in the adenomas of *Kras^wt/RR^* lungs, comparable to that of healthy airway epithelium. 4′,6-Diamidino-2-phenylindole (DAPI, blue) was used to visualize the cell nuclei. Scale bar: 50 µm. **c** Representative diagrams of FACS analysis of lung epithelial cells isolated from *Kras^wt/wt^* and *Kras^wt/RR^* mice 12 weeks after Ad-Flpo infection. Viable (Hoechst 33258-negative), CD31- and CD45-negative cells were visualized by epithelial cell adhesion molecule (EpCAM) expression and endogenous tdTomato fluorescence; tdTomato-positive cells (tdTomato^+^) are highlighted in the black frame, with their relative abundance indicated.

Trophoblast cell surface antigen 2 (Trop2) is a transmembrane glycoprotein with an as yet unknown physiological function. This antigen is highly expressed in most carcinomas, where it is associated with increased invasiveness and metastasis. In addition, Trop2 is expressed in the lungs of various organisms; in mice, Trop is mainly produced in the airway epithelium (Lenart et al. 2022). Our staining with an anti-Trop2 antibody confirmed this.

Neoplastic lesions that formed in the airways also showed positivity for Trop2, indicating that they were probably adenomas (Fig. 3B). We then investigated whether neoplastic cells carrying the recombined *Kras^RR^*allele could be isolated based on the red fluorescence of the tdTomato protein. Indeed, the “red” cells were only found in the cell suspension from the lungs of *Kras^wt/Flp-ready^* mice infected with Ad-Flpo, but not in control mice, i.e., the *Kras^wt/wt^* genotype (Fig. 3C).

### Kras^RR^ accelerates the growth of Apc-deficient tumors in the colon

Colon tumors, especially colorectal and rectal carcinomas, are among the most common solid tumors with an annual incidence of 1.7 million cases (Bray et al. 2018). Most of these tumors are triggered by the loss of tumor suppressor Apc, which leads to hyperproliferation of epithelial cells (Cancer Genome Atlas 2012). Oncogenic mutations in the *KRAS* gene are frequent mutational changes associated with the progression of adenomas after the loss of Apc (Dunne and Arends 2024). To investigate the functionality of the *Kras^RR^* allele in the context of intestinal tumors, we used a model with a conditional allele (cKO) of the *Apc* gene whose exon is flanked by two loxP sites. Cre-mediated recombination of this exon causes a frameshift resulting in a truncated and non-functional APC protein (Kuraguchi et al. 2006). In several of our previous studies, we used different transgenic mice, so-called Cre drivers, which allow targeted Apc deletion in different intestinal epithelial cells (Horazna et al. 2019; Hrckulak et al. 2018). In this case, we used rectal instillation of a Cre-producing adenovirus, Ad-Cre. Eight weeks after the administration of Ad-Cre, we examined the colonic epithelium and discovered dozens of relatively small proliferating lesions, hyperplastic crypts, which did not enlarge even 12 weeks after instillation. What is the basis for this phenotype? According to the conventional model of colorectal carcinogenesis, the tumor suppressor Apc must be inactivated in intestinal stem cells at the base of the intestinal crypts. One possible explanation is that the stem cells were not infected due to their location at the base of the crypts or due to the presence of a mucus barrier. The resulting lesions, which apparently cannot progress, occur after the loss of Apc in more differentiated cell types, as previously described by Barker and colleagues for tumors in the small intestine (Barker et al. 2009). Nevertheless, we and several other laboratories have documented that transgenic Cre-mediated Apc loss in intestinal stem cells leads to the formation of a larger number of neoplastic lesions in the small intestine, whereas relatively small hyperproliferative lesions that are not prone to progression form in the colon (Waaler et al. 2012). The phenotype described above is therefore more related to the particular biology of the mouse model and is not caused by the limited infectivity of the epithelial stem cells in the colon.

Although the role of Apc in intestinal tumors in humans is well documented (Vogelstein et al. 2013) and its tumorigenic role has been confirmed in mice (Stastna et al. 2019), some discrepancies between the human and mouse models have been identified. This mainly concerns the fact that adenoma formation in mice with Apc loss occurs predominantly in the small intestine and not in the large intestine, as is the case in human patients. It is hypothesized that the intestinal microbiota and diet play an important role (Dove et al. 1997). However, there is still no satisfactory explanation for this biological phenomenon.

After activation of the oncogenic *Kras^RR^* allele alone, we did not observe any hyperproliferative lesions or crypts. However, after administration of Ad-CreERT2-P2A-Flpo, we found large adenomas obstructing the colon 8 weeks after adenovirus administration, and the experimental animals had to be euthanized (Fig. 4A). A similar effect between Apc inactivation and oncogenic Kras activation was observed by Sansom and colleagues, who used floxed alleles of both genes and inducible Cre recombinase expression in Ah-Cre transgenic mice to inactivate the Apc allele and simultaneously induce the oncogenic *Kras* allele (Sansom et al. 2006). This suggests that, similar to human tumors, Kras^RR^ expression in transformed Apc-deficient cells causes tumor progression. The above results seem to contradict studies that have shown that the activation of Kras^G12D^ alone causes hyperproliferation of colonic epithelial cells. A major difference to our results is that the aforementioned studies used transgenic mice that produce a constitutively active, i.e., unregulated variant of Cre recombinase, which is controlled by the regulatory regions of genes that are active in most intestinal epithelial cells. The use of these Cre drivers led to continuous and sustained activation of the oncogenic Kras already in (late) embryonic development, which is probably related to the observed “hyperproliferative” phenotype (Calcagno et al. 2008; Haigis et al. 2008). Interestingly, it was previously shown that although the oncogenic Kras variant induces cellular hyperplasia not only in the colonic epithelium but also in other tissues, no morphologically detectable abnormalities were observed in the epithelium of the proximal part of the small intestine, regardless of the type of Cre driver used, whether regulated or unregulated (Haigis et al. 2008; Snippert et al. 2014).

**Figure 4.**
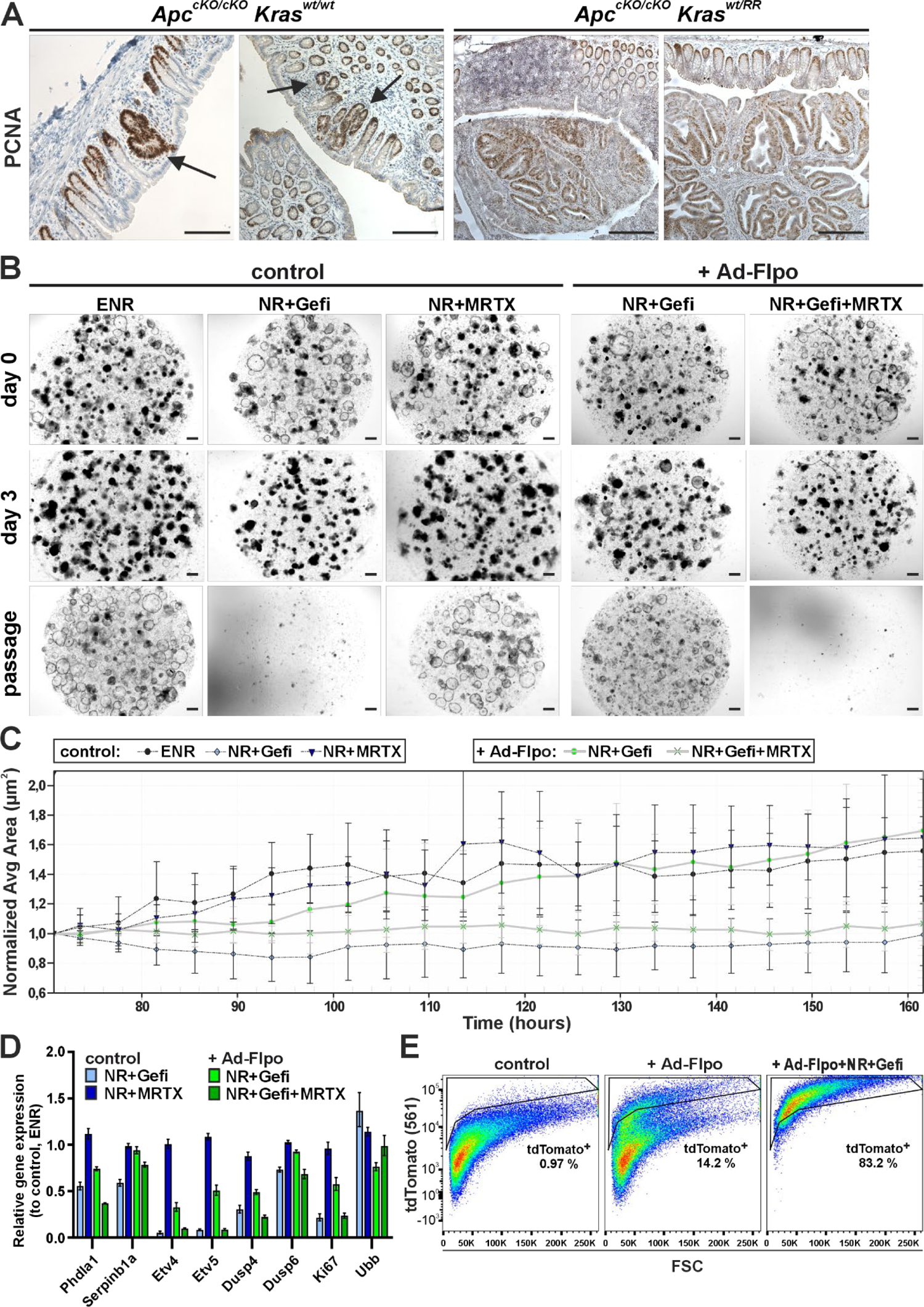
The influence of Kras^RR^ on the size of colon tumors and small intestinal organoids cultured with EGFR and Kras^G12D^ inhibitors **a** Immunostaining for proliferating cell nuclear antigen (PCNA, brown nuclei) in colon tissue sections from *Apc^cKO/cKO^/Kras^wt/wt^*and *Apc^cKO/cKO^/Kras^wt/RR^* mice, 10 weeks after intrarectal instillation of adenovirus particles expressing CreERT2 and Flpo (Ad CreERT2-P2A-FLPo). Hyperplastic lesions were present in the colon of *Apc^cKO/cKO^/Kras^wt/wt^*mice (black arrows), while large adenomas were detected in the colon of *Apc^cKO/cKO^/Kras^RR/wt^*mice. Sections were counterstained with hematoxylin (blue nuclei). Scale bar: 200 µm. **b** Growth of small intestinal organoids isolated from crypts of *Kras^wt/Flip-ready^* mice was monitored by brightfield microscopy after infection with Ad-Cre adenovirus particles (control) and Ad-FLPo particles (+ Ad-Flpo; right two columns). Organoids were cultured either in complete culture medium (ENR) supplemented with mouse EGF, noggin and R-Spondin1, or in the medium without EGF (NR) for Ad-Flpo-infected organoids. Seven days after infection, organoids were seeded and cultured for a further two days under the same conditions before addition of the EGFR tyrosine kinase inhibitor gefitinib (Gefi) or the Kras^G12D^ inhibitor MRTX1133 (MRTX). Shown are representative images of organoid cultures on the day the inhibitors were added (day 0), three days after the inhibitors were added (day 3) and two days after the first passage (passage). Scale bar: 200 µm. **c** Growth of control organoids and Kras^RR^-expressing organoids under different culture conditions was quantified by continuous *in vivo* imaging and automatic segmentation of organoid area (size). Control organoids cultured in ENR medium and Ad-Flpo-infected organoids cultured in NR medium with gefitinib were each seeded in eight technical replicates. Seventy-two hours after seeding, MRTX1133 was added to the culture medium. The average relative change in organoid size was recorded every 4 hours for the following 92 hours. While MRTX1133 did not affect the growth of control organoids, gefitinib led to a growth arrest. Conversely, gefitinib had no effect on the growth of Kras^RR^-expressing organoids when used alone, but the addition of MRTX1133 completely inhibited the growth of Ad-Flpo-infected organoids. The error bars represent the standard deviation, and the average organoid area values were normalized to the time of application of the inhibitors. **d** Reverse transcription and quantitative PCR analysis was used to compare the expression of common MAPK signaling pathways and proliferation markers in *Kras^wt/Flip-ready^* small intestinal organoids infected with adenoviral control particles (control; blue columns) and Ad-Flpo particles (+ Ad-Flpo; green columns). The control organoids were expanded in ENR medium, the Ad-Flpo-infected organoids in NR medium with gefitinib. The graph shows the log2 fold change in expression of selected genes in organoids cultured in different media compared to their expression in Ad-Cre-infected organoids cultured in the ENR medium. Total RNA was isolated 48 hours after application of the selective conditions. The relative expression of β-actin was arbitrarily set to 1; the expression of housekeeping gene ubiquitin B (*Ubb*) is also shown. The experiment was performed in technical triplicates; the error bars indicate the confidence interval. **e** Representative graphs show FACS analyses of cells isolated from *Kras^wt/Flip-ready^*organoids of the small intestine infected with adenoviral control particles (left) or Ad-Flpo particles (center and right) and cultured in ENR or selective medium (NR+Gefi). Viable (Hoechst 33258-negative) cells were visualized by endogenous tdTomato fluorescence and forward scatter (FSC); tdTomato^+^ cells are shown in the black frame with their relative abundance. Control cells (left) showed minimal background tdTomato fluorescence, while a discrete population of Kras^RR^-expressing cells appeared after Ad-Flpo infection (center). These tdTomato^+^ cells were further enriched after growth under selective conditions (NR+Gefi; right), while non-recombined cells were lost during two passages.

To study the effects of activation of the *Kras^RR^* allele alone (by Flpo recombinase) and the effects of simultaneous inactivation of the *Apc* gene, we used an adenoviral vector carrying the gene for tamoxifen-regulated Cre fused to a modified version of the estrogen receptor domain (ERT2). After the *CreERT2* gene, the sequence for peptide 2A (*P2A*) of porcine teschovirus-1 and the gene encoding Flpo were inserted. However, testing of this adenoviral vector with MEFs from mice containing the Cre-dependent reporter allele *R26-tdTomato* and the Flp-dependent reporter allele *R26-Blueflirt* revealed that tamoxifen regulation of the CreERT2 protein was not functional (data not shown). This effect may be related to the impaired functionality of the ERT2 domain in the context of the fusion protein. Although it has been documented that cleavage of fusion proteins at the P2A site is highly efficient, some of the proteins containing P2A in the form of an uncleaved precursor remain in the cell (Provost et al. 2007), which may result in a loss of ERT domain-mediated regulation of Cre localization and nuclear activity. Another possibility is that Cre regulation by the ERT domain is not 100% effective, and “leaky” enzyme activity has been documented even in the absence of tamoxifen (Kristianto et al. 2017).

### Analysis of the effects of the *Kras^RR^* allele on intestinal organoids

To analyze the effects of Kras^RR^ expression on intestinal epithelial cells, we used organoids prepared from the small intestine of *Kras^wt/Flp-ready^* mice. First, we investigated whether the organoids can grow *in vitro* after adenoviral recombination independently of epidermal growth factor. Since the control organoids can survive without exogenous Egf (Sato et al. 2011), we added the Egfr inhibitor gefitinib (Kris et al. 2003) to the medium. We were able to show that gefitinib stopped the growth of control organoids infected with Ad-Cre and did not allow the culture to survive passage. After recombination with Ad-Flpo, the growth of organoids in the medium with gefitinib slowed down for one passage, but then they continued to grow indefinitely (Fig. 4 B and C). The initial slowdown was due to the ongoing selection of successfully recombined cells expressing Kras^RR^.

We then investigated whether the growth ability of mutant organoids was impaired under Egfr inhibition by the recently discovered KRAS^G12D^-specific inhibitor MRTX1133 (Hallin et al. 2022). The growth of control organoids showed that MRTX1133 had no effect on cells with wt Kras. In contrast, when MRTX1133 was added to Kras^RR^-positive organoids selected by the presence of gefitinib, the growth of the culture stopped and the organoids did not survive passage (Fig. 4, B and C). Furthermore, expression analysis of target genes in the organoids two days after addition of the inhibitor confirmed that the effect of Kras^RR^ on the survival and proliferation of intestinal epithelial cells is mediated by MAPK signaling. While the expression of target genes of the MAPK signaling pathway and proliferation marker Ki67 was reduced in control organoids after the addition of gefitinib, similar levels were only achieved in Kras^RR^-positive organoids after the addition of MRTX1133 (Fig 4D). Of note, the expression of MAPK pathway target genes and Ki67 was lower in Kras^RR^-positive organoids during gefitinib selection. As mentioned above, this may be due to the presence of a certain amount of non-recombined cells in the early phase of selection. However, we cannot exclude the possibility that the dose of the active Kras form in cells with blocked EGFR receptor plays a role. MRTX1133 had no effect on the expression level of the analyzed genes in control cells (Fig. 4D). The selection process of Kras^RR^-positive cells by gefitinib during two passages after recombination *in vitro* was recorded by flow cytometry and showed an enrichment of tdTomato-positive cells after growth under the selective conditions (Fig. 4 E).

The RAS family proteins have an extraordinarily high affinity for GTP, and it seemed almost impossible to find a compound that could block GTP-RAS binding (Ledford 2015). However, in 2013, Ostrem and colleagues described a compound that blocks the function of mutant Kras protein KRAS^G12C^ by irreversibly binding to cysteine at position 12 (Ostrem et al. 2013). Currently, selective KRAS^G12C^ inhibitors, which occur in about 20% of lung cancer cases and 3% of colorectal cancer cases, are being tested in clinical trials for various human tumor types (Fakih et al. 2022; Yaeger et al. 2023). The molecule MRTX1133 is an example of a potent selective non-covalent KRAS^G12D^ inhibitor. Given the fact that KRAS^G12D^ is three times more common in human neoplasms than the KRAS^G12C^ variant, the clinical importance of KRAS^G12D^ inhibitors is obvious. Furthermore, the development of panKRAS inhibitors that are also effective against more frequent mutation variants such as G12D, G12V, G12A, and G13C/D in addition to G12C is very promising. One such inhibitor is currently undergoing preclinical testing and shows great potential (Kim et al. 2023).

In summary, the *Kras^RR^* allele, which allows analysis (sorting) of cells producing the KrasG12D oncogene based on red fluorescence, could be an important tool to evaluate the specificity, mechanism of action, and effects of short and long-term administration of inhibitors of oncogenic KRAS variants. This could be done *in vitro* using organoids or *in vivo* in combination with a suitable transgenic strain that produces Flpo recombinase in different tissues.

## Abbreviations

APC: adenomatous polyposis coli

cKO: conditional allele

E: embryonic day

EGF: epidermal growth factor

EGFR: epidermal growth factor receptor

ERT: estrogen receptor ligand-binding domain

ES cells: embryonic stem cells

Flp: flippase

Flpo: codon-optimized version flippase

FRT: Flp recombinase recognition site

GAP: GTPase-activating protein

KRAS: KRAS proto-oncogene

GTPase Kras^RR^: RedRas allele producing Kras^G12D^

tdTomato; MAPK: mitogen-activated protein kinase

MEFs: mouse embryonic fibroblasts

RT-qPCR: reverse transcription quantitative PCR

Trop2: trophoblast cell surface antigen 2

wt: wild-type

## Acknowledgement

We thank S. Takacova for critically reading the manuscript. This work was supported by the he Czech Science Foundation, grant no. 20-31322S, and by the project National Institute for Cancer Research (Program EXCELES, ID Project No. LX22NPO5102, Funded by the European Union - Next Generation EU). We thank the staff of the Transgenic unit for blastocyst injections, Zdenek Cimburek for technical assistance, and Eva Sloncova and Katerina Galuskova for their administrative and technical support.

